# The Fluorescent Dye 1,6-Diphenyl-1,3,5-Hexatriene Binds to Amyloid Fibrils Formed by Human Amylin and Provides a New Probe of Amylin Amyloid Kinetics

**DOI:** 10.1101/2021.05.10.443442

**Authors:** Ming-Hao Li, Lakshan Manathunga, Erwin London, Daniel P. Raleigh

## Abstract

The fluorescent dye 1,6-diphenyl-1,3,5-hexatriene (DPH) is widely used as a probe of membrane order. We show that DPH also interacts with amyloid fibrils formed by human amylin (also known as islet amyloid polypeptide) in solution and this results in a 100-fold increase in DPH fluorescence for a sample of 20μM human amylin and 0.25 μM DPH. No increase in DPH fluorescence is observed with the non-amyloidogenic rat amylin or with freshly dissolved, non-fibrillar human amylin. The time course of amyloid formation by amylin was followed by monitoring the fluorescence of added DPH as a function of time and was similar to that monitored by the standard fluorescent probe thioflavin-T. The inclusion of DPH in the buffer did not perturb the time course of amyloid formation under the conditions examined and the time course was independent of the range of DPH concentrations tested (0.25 to 5 μM). Maximum final fluorescence intensity is observed at substoichiometric ratios of DPH to amylin. No significant increase in fluorescence was observed during the lag phase of amyloid formation, and the implications for the structure of amylin pre-fibril oligomers are discussed. Human amylin contains three aromatic residues. A triple aromatic to leucine mutant forms amyloid and DPH binds to the resulting fibrils, indicating that interactions with aromatic side chains are not required for DPH amylin amyloid interactions. DPH may be especially useful for studies on mutant amylins and other polypeptides in which changes in charged residues might complicate interpretation of thioflavin-T fluorescence.

## Introduction

Amyloid formation refers to the aggregation of proteins and polypeptides into partially ordered fibril structures rich in cross-β structure. The process of amyloid formation has been implicated in more than 30 different human diseases, and a large number of proteins which do not form amyloid *in vivo* can be induced to do so *in vitro*^*1-3*^. Prominent examples of proteins involved in disease related amyloids include the Aβ peptide and tau protein of Alzheimer’s disease, alpha-synuclein in Parkinson’s disease and amylin, (also known as islet amyloid polypeptide, IAPP) in type-2 diabetes^*1-5*^. Amyloid formation is commonly monitored *in vitro* using fluorescence-based dye binding assays in which the added dye displays an increase in fluorescence when bound to fibrils. Probably the most widely used fluorescence amyloid probe is the cationic benzylamine-benzothiazole based dye, thioflavin-T (Figure-1). Thioflavin-T undergoes a significant increase in fluorescence upon binding to amyloid fibrils and usually does not exhibit enhanced fluorescence in the presence of pre-amyloid species or monomeric proteins^6, 7^. There is interest in the discovery and development of new amyloid binding dyes since thioflavin-T, and other dye-based assays of amyloid formation can give both false positives and false negatives^8-13^. The cationic nature of thioflavin-T also indicates that it may not be optimum for following amyloid formation by polypeptides which have a significant net positive charge^14^. In some cases, dye binding studies can provide information about the properties of amyloid fibrils or pre amyloid intermediates^15-17^.

The fluorescence of 1,6 diphenyl 1,3,5 hexatriene (DPH, Figure-1) is weak in water, but increases significantly in a hydrophobic environment^18, 19^. The dye is widely used as a probe of membrane order, an assay that relies on changes in its fluorescence anisotropy ^19-21^. It can also be used to detect the formation of detergent micelles or detect the level of lipid vesicles by the enhancement of its fluorescence intensity in the presence of a hydrophobic aggregate^22,23^. Here we demonstrate that the fluorescence of DPH is also enhanced in the presence of amyloid fibrils formed by human amylin (h-amylin, also known as islet amyloid polypeptide or IAPP). The probe can be used to accurately follow the kinetics of h-amylin amyloid formation, but shows no significant fluorescence enhancement in the presence of the non-amyloidogenic variant of amylin derived from rats and mice or in the presence of freshly dissolved non-fibril h-amylin.

Islet amyloid formation by h-amylin contributes to pancreatic β-cell dysfunction and death in type 2 diabetes (T2D), and in islet transplants^5, 24-33^. h-Amylin is a cationic 37 residue neuro pancreatic polypeptide which has a conserved amidated C-terminus and a conserved disulfide bridge between residues 2 and 7 (Figure-1). The polypeptide lacks acidic residues and has a net charge at physiological pH of between +2 and +4, depending on the pKa of the N-terminus and His-18. h-Amylin is one of the most amyloidogenic naturally occurring sequences known and forms amyloid *in vitro* more rapidly than the Aβ peptide of Alzheimer’s disease ^28^. The sequence of h-amylin is displayed in figure-1 together with the structures of DPH and thioflavin -T. There are limited investigations of the use of DPH to follow amyloid formation. Prior studies have shown that DPH can bind to aggregates of Aβ and α-synuclein, but its ability to monitor amylin formation has not been tested and its suitability as a dye for *in situ* kinetics assays has not been investigated in depth^34^.

**Figure-1:**
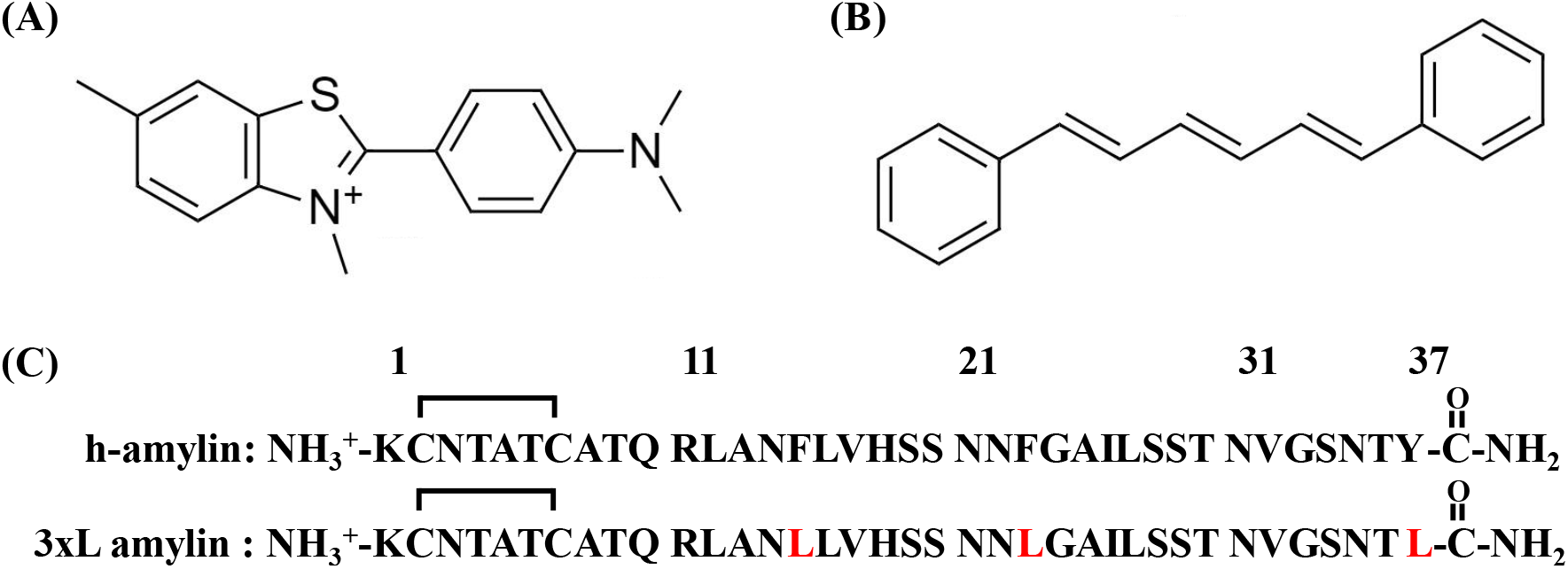
The structure of (A) Thioflavin-T and (B) 1,6 diphenylhexatriene. (C) The primary sequences of h-amylin and the triple aromatic to Leu mutant of h-amylin. The polypeptides have an amidated C-terminus and an intramolecular disulfide between Cys-2 and Cys-7. Residues in the triple mutant which differ from wild type are in red.

## Results and Discussion

### DPH exhibits enhanced fluorescence in the presence of h-amylin amyloid fibrils, but not in the presence of non-amyloidogenic forms of h-amylin or rat amylin

We formed amyloid fibrils of h-amylin by incubating the polypeptide in buffer at pH 7.4. Fibril formation was verified by thioflavin-T assays and by transmission electron microscopy (TEM). DPH exhibited a significant increase in fluorescence with an emission maximum of 427 nm when added to the pre-formed h-amylin fibrils. In contrast, no significant DPH fluorescence was detected in the presence of freshly dissolved, no n-fibril h-amylin (Figure-2). Not all variants of amylin form amyloid, and mouse/rat amylin is not amyloidogenic *in vitro* under standard conditions, nor do rats or mice develop islet amyloid or diabetes^4, 26^. DPH showed no increase in fluorescence in the presence of rat amylin (Figure-2).

**Figure-2:**
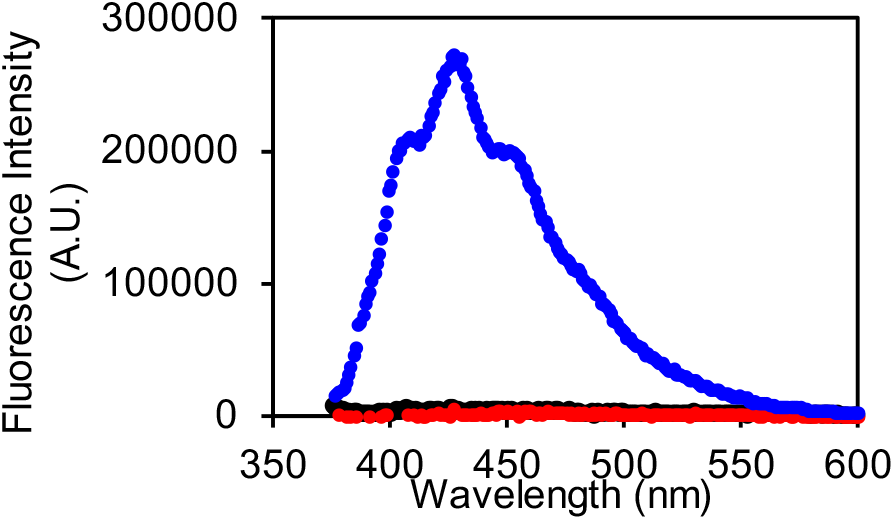
Fluorescence emission spectra of DPH in the presence of h-amylin fibrils (blue), freshly dissolved non-fibril h-amylin (black), and rat amylin (red). The black and red curves overlap. Experiments were conducted with 0.25 μM DPH, at pH 7.4 25 °C in 20 mM tris 100 mM NaCl buffer. The concentration of rat and human amylin were 20 μM.

### DPH fluorescence accurately reports on the kinetics of h-amylin formation

Fluorescence based dye binding assays are widely used to study the kinetics of amyloid formation and to detect amyloid fibrils^6, 7^. They are most convenient when the dye is added at the start of an assay and its fluorescence continuously monitored. However, dyes are extrinsic probes, and the possibility always exists that they could perturb the kinetics of the process they are designed to monitor. Dyes are still useful in this case, but a kinetic experiment will then need to be conducted by removing aliquots of a reaction mixture at different times points, adding dye, and measuring the fluorescence signal. This is much more time consuming than *in situ* experiments and does not lead itself to high throughput experiments. Consequently, it is important to test if a dye perturbs the time course of amyloid formation.

Amyloid formation by h-amylin displays the typical kinetic progress curve observed for amyloidogenic proteins; a lag time in which few if any detectable amyloid fibrils are formed, followed by a growth phase which results in increased material deposited as fibrils, and increased dye fluorescence. The system reaches a plateau phase in which fibrils and any remaining soluble polypeptide is in equilibrium. This leads to a sigmoidal curve of fluorescence vs time. Thioflavin-T has previously been showed to not perturb the kinetics of amyloid formation by h-amylin under the conditions of our experiments^5, 17, 32^. Thus, we conducted side by side kinetic experiments with h-amylin to test if *in situ* experiments with DPH accurately report on h-amylin amyloid formation. One set of experiments was run with added thioflavin-T as the reporter dye and another set used DPH as the probe. Initial experiments indicated that DPH had a tendency to bind to the standard plastic 96 well plates (Corning non-binding plates) used in our plate reader fluorescence assays. We did not attempt to find optimum plastic plates for the DPH assays, but instead used a quartz plate for the proof-of-principle studies reported here. The time courses detected by DPH and thioflavin-T are similar and give the same value of T_50_ (9 h) the time required to reach 50% of the final intensity in an assay (Figure-3). This indicates that DPH is as reliable at monitoring h-amylin amyloid formation as thioflavin-T. A 100-fold increase in DPH fluorescence was observed when a 20 μM sample of h-amylin formed amyloid fibrils in the presence of 0.25 μM DPH dye, and an even larger enhancement is observed for higher DPH concentrat ions (Fiugre-4). For comparison, 20 μM thioflavin-T lead to a 100-fold fluorescence when a 20 μM sample of h-amylin formed amyloid fibrils.

We further tested the use of DPH in *in situ* studies by conducting a set of experiments in which the concentration of added DPH spanned a 20-fold range, from 0.25 μM to 5 μM for a fixed concentration of h-amylin (40 μM). If DPH modulated the time course of amyloid formation, then one would expect more pronounced effects as the concentration of the dye was increased. No detectable DPH concentration dependent effects were observed, providing additional evidence that the dye does not alter the time course of amyloid formation (Supporting Figure-1).

We next examined the effect of varying DPH concentration on the final fluorescence observed in the plateau region after amyloid formation is complete. The final DPH fluorescence intensity appeared to saturate above 2 μM DPH for 20 μM h-amylin under these conditions (Figure-4).

**Figure-3:**
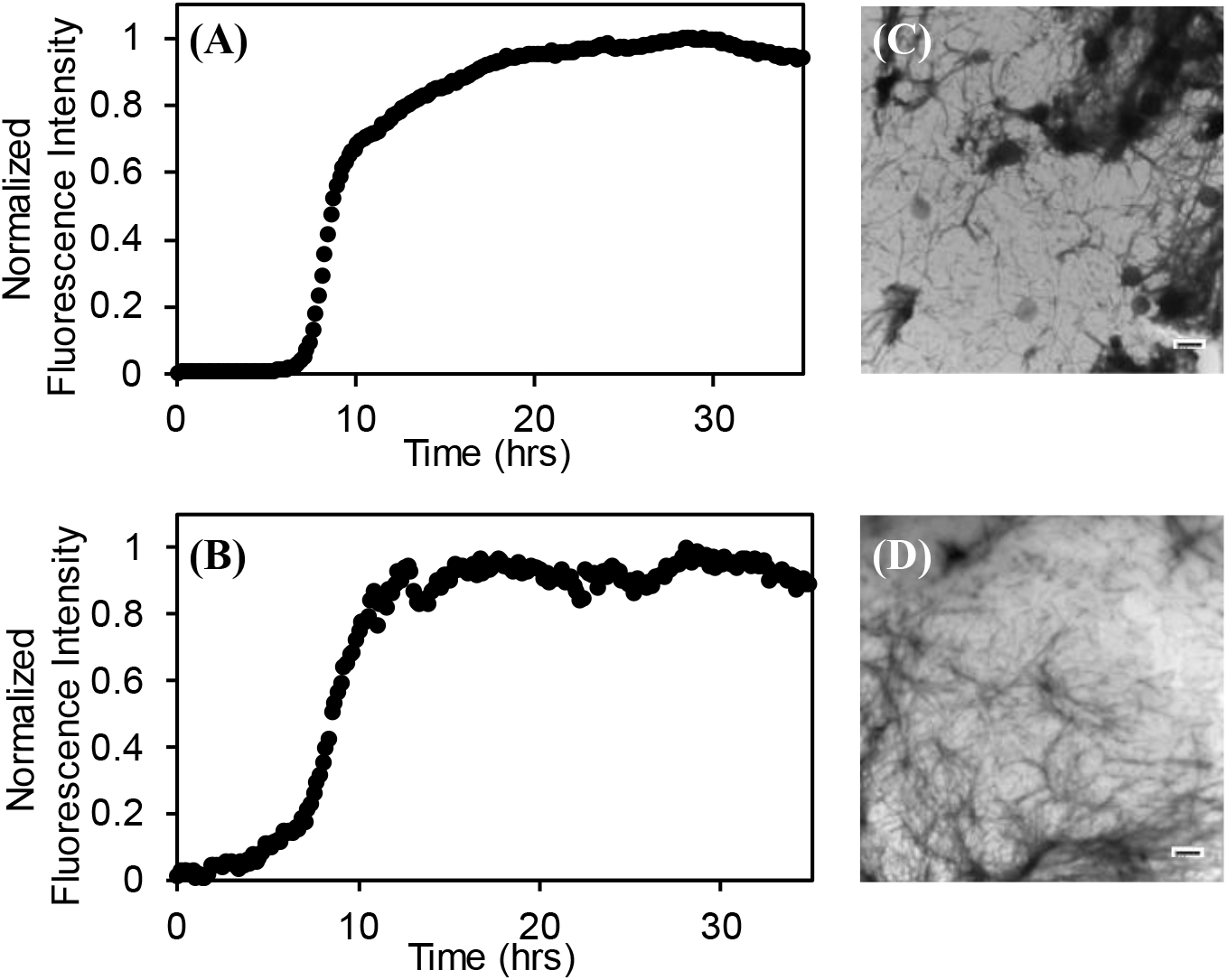
The kinetics of h-amylin amyloid formation followed by (A) a thioflavin-T assay with a 40 μM thioflavin-T concentration and (B) a DPH assay with a 0.25 μM DPH concentration. The signal to noise of the DPH experimental can be enhanced significantly by increase the DPH concentration. TEM images of samples collected at the end of the (C) thioflavin-T assay and the (D) DPH assay. Experiments were conducted at 25 °C, pH 7.4 in 20 mM tris 100 mM buffer. The concentration of h-amylin was 20 μM in both experiments and the concentration of the dyes were 40 μM for thioflavin-T and 0.25 μM for DPH.

**Figure-4:**
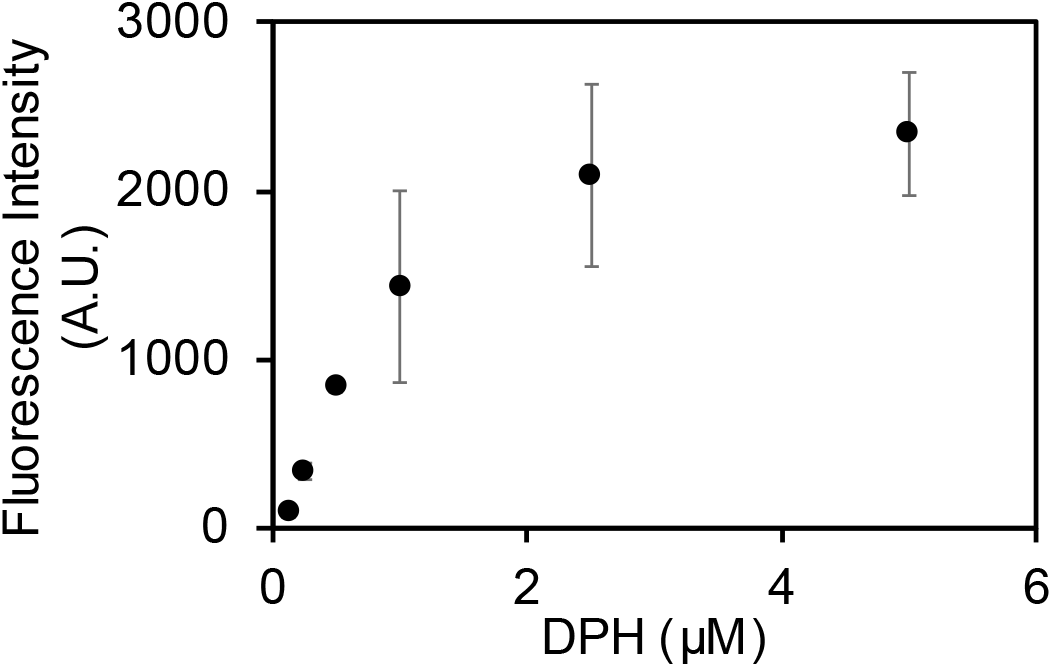
A plot of the final intensity observed in a set of DPH assays. Assays were conducted in a 96 well quartz plate using a plate reader at 25 °C, pH 7.4 in 20 mM tris 100 mM buffer. The concentration of h-amylin was 20 μM.

### π-π interactions are not required for DPH h-amylin fibril interactions

h-Amylin contains three aromatic residues, two Phe residues at positions 15 and 23 and a strongly conserved Tyr at the C-terminus (Figure-1, Figure-5). Given the highly conjugated structure of DPH, we wondered if the aromatic residues in h-amylin play a critical role in binding DPH via π-π interactions. There are several structural models of the h-amylin fibrils formed *in vitro*. One based on solid state NMR studies, one on crystallographic studies of small steric zipper-forming peptides derived from amylin, and three cryo-EM structures^33, 35-38^. The different structures differ in their details and this may be due to the fact that the fibrils were formed under different conditions, but all contain extensive cross β-sheet structure, and the fibril is made up of two stacks of h-amylin monomers (Figure-5).

Each model contains several aromatic residues within the cross-β core; this leads to a quasi-infinite arrangement of stacked aromatic sidechains. Analysis of the solvent exposed surface area reveals different levels of exposure of the aromatic residues in the different models. In two of the three cryo-EM based models, all three aromatics are largely buried. The % exposed solvent accessible surface area (% SASA) is less than 4% for all residues in both of the stacks in 6Y1A and 6ZRF. In 6VW2, F15 and Y37 have 53 to 84 % and 40 to 53% SASA respectively. The quoted range reflects for the values in the two different stacks that make up the fibril structure. In the model based on x-ray studies of steric zipper peptides, F15 and F23 exhibit 38 to 60 % and 37 to 59% SASA, respectively. In the model based on solid state NMR studies F23 is largely buried (% SASA from 10 to 17%), but the % SASA of F15 and Y37 range from 47 to 74% and 40 to 74%. We tested whether the aromatic residues are required for DPH binding to amylin amyloid fibrils. A triple aromatic to Leu mutant of h-amylin (F15L, F23L, Y37L, amylin, denoted as 3xL-amylin) has been shown to form amyloid, albeit it at a slower rate than wild type^5, 17, 32^. There is a significant increase in DPH fluorescence when the dye is added to 3xL amylin fibrils, but not when it is added to freshly dissolved non fibril 3xL amylin (Figure-6, supporting Figure-S2). The intensity at the fluorescence emission maximum is somewhat less for the 3xL amylin variant than for the wild type. This could arise because the binding of DPH is weaker, its quantum yield when bound is less, or because fewer fibrils are for med by the mutant. These experiments demonstrate that aromatic residues are not required for DPH to interact with amylin amyloid fibrils.

**Figure-5:**
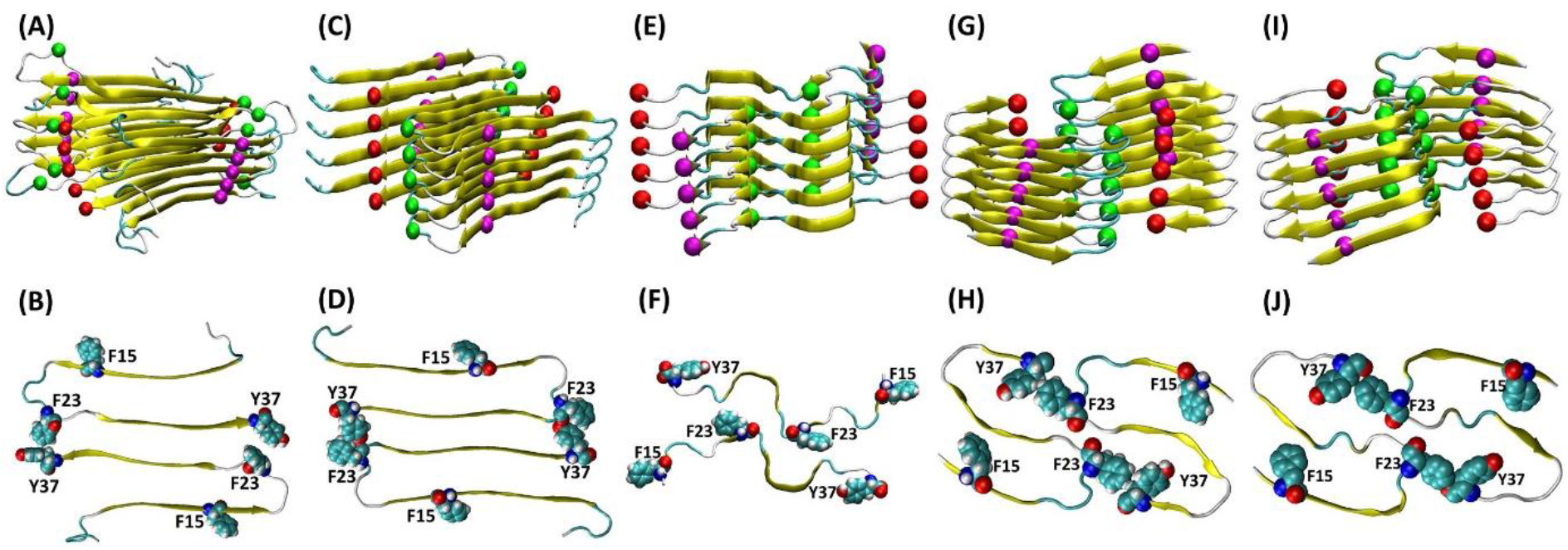
Structural models of the h-amylin amyloid fibril with the location of the aromatic residues shown. The top row displays ribbon diagrams showing the arrangement of individual polypeptides in the fibril and spheres showing the Cα atoms of Phe-15 (magenta), Phe-23 (green) and Tyr-37 (red). The bottom row shows a top-down view and displays a cross section of the fibril with the aromatic residues shown in space-filling format. (A, B) structure based on solid state NMR studies. (C, D) structural model based on x-ray diffraction studies of small steric zipper peptides. (E, F) Cryo-EM structure of a h-amylin fusion determined at pH 8.0 (pdb code 6VW2). (G, H) Cryo-EM structure determined at pH 6.0 (pdb code 6Y1A). (I, J) Cryo-EM structure determined at pH 6.8 (pdb code 6ZRF).

**Figure-6:**
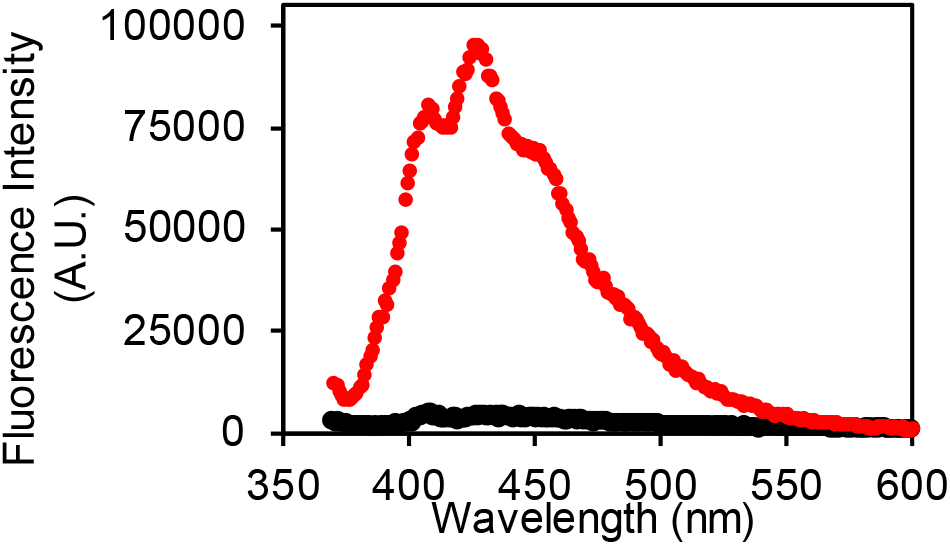
Fluorescence emission spectra of DPH in the presence of: 3xL-amylin fibrils (red), and freshly dissolved non-fibril 3xL-amylin (black). Experiments were conducted with 20 μM peptide, 0.25 μM DPH, at pH 7.4, 25 °C in 20 mM tris 100 mM NaCl buffer.

## Conclusions

The features of the fibril that lead to DPH binding are not known, but the cross β-structure leads to long exposed faces with grooves on the surface of the fibril. Thioflavin-T is believed to bind in these groves and DPH may as well. The studies reported here show that DPH accurately reports on the kinetics of amyloid formation by amylin and demonstrates that interactions between aromatic residues and DPH are not required for DPH binding. The dye may be used at sub-stoichiometric concentrations, indeed some of the experiments reported here involved an 80-fold excess of h-amylin to DPH. DPH may be especially useful for studies on mutant amylins and other polypeptides in which changes in charged residues might complicate interpretation of thioflavin-T fluorescence.

Dye binding studies have been used to probe the development of hydrophobic surfaces/ clusters in pre-fibril oligomers. In particular, some amyloidogenic proteins have been reported to bind dyes such as ANS, bis-ANS, and Nile Red in the lag phase and ANS binding has been suggested to be a general property of oligomers formed by a wide range of proteins^15, 16^. Thus, it is interesting to note that no significant enhancement of DPH fluorescence was observed in the lag phase of h-amylin amyloid formation. This is consistent with previous studies which have shown that h-amylin lag phase species do not bind ANS or Nile Red^17^ and argues that h-amylin oligomers do not present large well-developed solvent exposed hydrophobic surfaces.

## Materials and Methods

### Peptide Synthesis and Purification

h-Amylin and the triple aromatic to Leu mutant were prepared by solid phase peptide synthesis on a 0.01 mmol scale with 9-fluorenylmethyloxycarbonyl (Fmoc) protected amino acids. The synthesis was conducted using a CEM Liberty microwave peptide synthesizer. Pseudoproline dipeptide (oxazolidine) derivatives of Leu-Ser were used as previously described to improve the yield^39,40^. 5-(4’-Fmoc-aminomethyl-3’,5’-dimethoxyphenol) valeric acid resin was employed. This resin provides an amidated C-terminus upon cleavage. Coupling reactions were carried out for 2 min at 90 °C except for Cys and His which were coupled at 55 °C to minimize the risk of racemization. Peptides were cleaved using a cocktail made of 92.5% trifluoroacetic acid (TFA), 2.5% water, 2.5% Triisopropyl silane (TIPS), and 2.5% 3,6-dioxa-1,8-octanedithiol (DODT). Cleaved crude peptides were dissolved in 20% acetic acid (v/v) and lyophilized prior to the formation of the disulfide bond. The intramolecular disulfide bond was formed by dissolving the dry peptide in 100 % dimethyl sulfoxide (DMSO) at a concentration of 10 mg/mL and incubating at room temperature for 3 days with stirring^41^. Both polypeptides were purified via reverse-phase high pressure liquid chromatography (HPLC) with a C18 preparatory column (Higgins Analytical) using a binary gradient (buffer A =100% H_2_O and 0.045% HCl and buffer B = 80% Acetonitrile, 20% H_2_O, and 0.045% HCl). HCl was used as a counter ion instead of TFA since TFA can affect amylin kinetic assays. Residual scavengers were removed after the first round of HPLC purification using a 1,1,1,3,3,3-hexafluoro-2-propanol (HFIP) extraction procedure. Peptides, after the first HPLC purification were dissolved in neat HFIP and incubated at room temperature for 4 h, then filtered with 0.45 μM GHP membrane filter, and purified via RP-HPLC. Masses of purified peptides were confirmed using matrix assisted laser desorption time-of-flight mass spectrometry; h-amylin expected: 3903.3, observed: 3902.7; 3xL-amylin expected: 3785.3 observed: 3785.9.

### Preparation of Peptide Stock Solutions and Sample Preparation

Stock solutions were prepared by dissolving the purified peptides into neat HFIP to a concentration of 1.6 mM (determined by lyophilizing a 20 μL aliquot of the HFIP stock for at least 12 h, dissolving in 120 μL of 20 mM Tris buffer (pH 7.4) and measuring the absorbance at 280 nm for h-amylin). An extinction coefficient of 16 15 M^-1^cm^-1^ at 280 nm was used for h-amylin. The concentration of the triple Leu mutant was determined using the BCA assay.

### DPH Fluorescence Emission Spectra

Fluorescence emission spectra were recorded in a 1 cm pathlength cell on a Horiba Quanta Master Spectrofluorimeter (Horiba Scientific, Edison, NJ). The excitation wavelength was 358 nm, and the excitation and emission slits were 4 nm. Spectra were recorded over the range 370 to 600 nm at 25 °C.

### DPH and Thioflavin-T Fluorescence Based Kinetic Assays

Aliquots of the HFIP stock solutions were lyophilized for at least 12 h and reconstituted in buffer at pH 7.4 for kinetic assays. The final concentrations of the peptides and dye (DPH or thioflavin-T) were 20 μM and 0.25 μM, respectively. A quartz plate was used for the kinetic assays. Wells at the edge of the plate were not used but were filled with buffer. Fluorescence assays were conducted using a Molecular Devices SpectraMax Gemini EM microplate reader. Fluorescence was measured from the bottom of the wells at 10 min intervals without agitation. The excitation and emission wavelengths for the DPH studies were 358 and 427 nm, respectively. The values for the thioflavin-T experiments were 450 nm (excitation) and 485 nm (emission). Three parallel samples were tested for each experiment. The quoted uncertainties are the estimated standard deviations. All fluorescence assays were performed at 25 °C.

### Transmission Electron Microscopy

Samples for TEM were prepared from the samples at the end of the kinetics experiments. 15 μL of peptide solution was blotted on a carbon-coated Formvar 300-mesh copper grid for 1 min and then negatively stained with 2% (w/v) uranyl acetate for 1 min. A FEI Bio TwinG^2^ transmission electron microscope was used to record TEM images (Life Science Microscopy Center at Stony Brook University).

### Calculation of Solvent Accessible Surface Area of the Aromatic Residues

The values of the SASA were calculated for each aromatic residue using the coordinate files with hydrogens included. The “measure SASA” command in Visual Dynamics Software (VMD)^42^ was used with a probe radius of 1.4 Å for the calculations. Values are reported as % relative to an extended tripeptide of the same sequence for F15 and F23 and relative to a C-terminal capped Thr-Tyr for the c-terminal tyrosine (Y37). Tripeptides and the capped dipeptide were generated using TLEAP in Amber20^43^.

### Associated Content

#### Supporting Information

One figure showing kinetic curves for a set of DPH monitored amyloid assays with a range of DPH concentrations. One figure showing TEM images of the amyloid fibrils formed by 3xL amylin in the presence of thioflavin-T and DPH.

The Supporting Information is available free of charge at: xxx

## Abbreviations

DODT: 3,6-dioxa-1,8-octanedithiol
DPH: 1,6-diphenyl-1,3,5-hexatriene
Fmoc: 9-fluorenylmethyloxycarbonyl
h: hours
h-amylin: human amylin
HFIP: 1,1,1,3,3,3-hexafluoro-2-propanol
HPLC: high performance liquid chromatography
IAPP: islet amyloid polypeptide
min: minutes
SASA: solvent accessible surface area
T2D: type-2 diabetes
T_50_: the time required to reach 50% of the final maximal fluorescence changes in a kinetic experiment
TEM: transmission electron microscopy
TFA: trifluoroacetic acid
TIPS: Triisopropyl silane
3xL-amylin: a triple F15L, F23L, Y37L mutant of h-amylin

## Author Contributions

MHL and LM conducted experiments, prepared regains, and analyzed data. DPR and EH analyzed data, supervised, and directed the research. The manuscript was written through the contributions of all authors. All authors have given approval to the final version of the manuscript.

## Acknowledgments

We thank Dr. Rehana Akter and Ms. Daeun Noh for helpful discussions concerning amyloid specific dyes. This work was supported by NIH grants GM 078114 and GM 122493 and by NSF grant MCB-1330259.

