## Supplemental Figure for "The Fluorescent Dye 1,6-Diphenyl-1,3,5-Hexatriene Binds to Amyloid Fibrils Formed by Human Amylin and Provides a New Probe of Amylin Amyloid Kinetics"

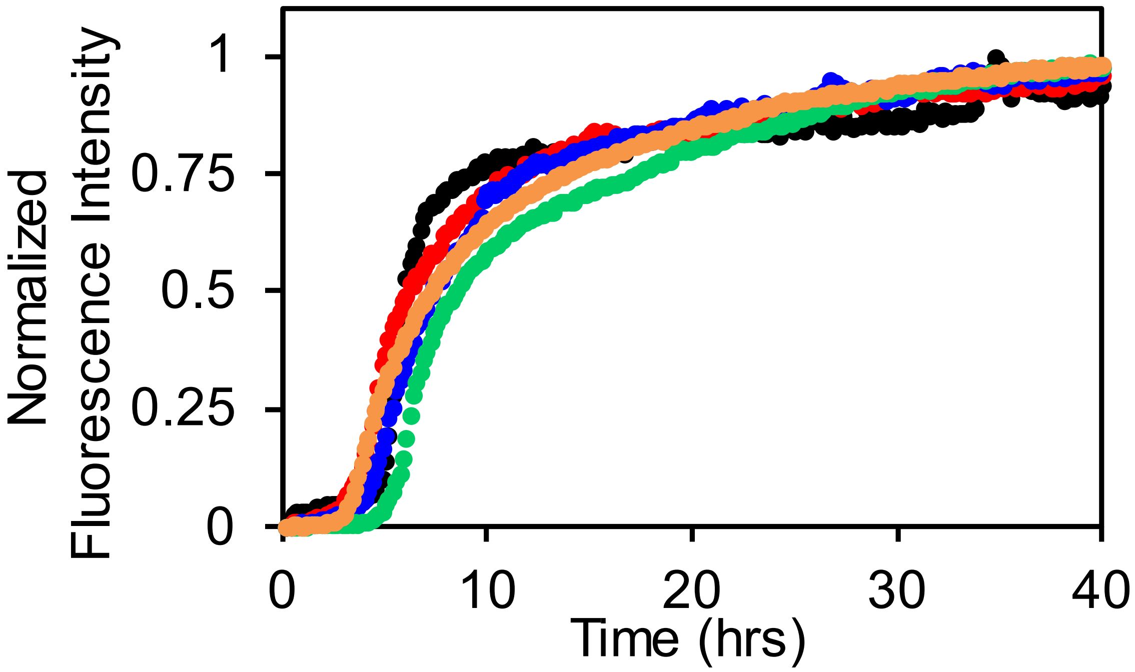


**Supporting Figure S1:** A comparison of the time course of h-amylin amyloid formation monitored using different concentrations of DPH. The vertical axis is plotted on a normalized scale. 0.25 mM (black), 0.5 mM (red), 1 mM (blue), 2 mM (green), and 5 mM DPH (orange). The concentration of h-amylin was 40 μM at pH 7.4 in 20 mM tris 100 mM NaCl buffer 25 °C for both experiments.

(A) (B)

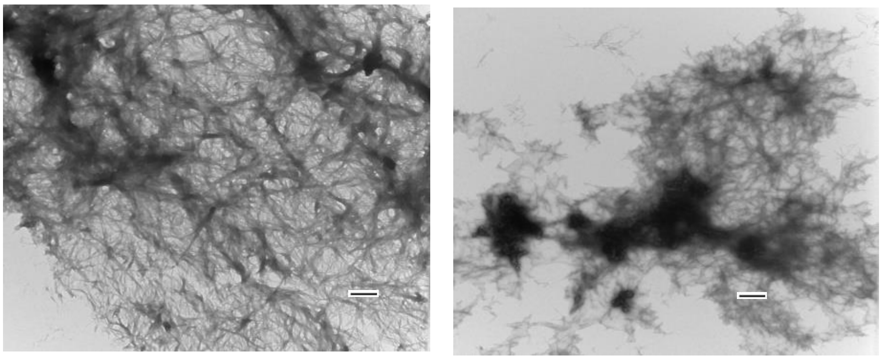

**Supporting Figure S2:** TEM images of samples collected at the end of the thioflavin-T and DPH assays for the triple Leucine mutant are displayed. (A) A sample of 20 μM of 3xL-amylin and 40 μM of thioflavin-T, and (B) a sample of 20 μM of 3xL-amylin and 0.25 μM of DPH at pH 7.4 in 20 mM tris 100 mM NaCl buffer. Scale bars represent 100 nm.
